# TFBindFormer: A Cross-Attention Transformer for Transcription Factor–DNA Binding Prediction

**DOI:** 10.64898/2026.04.09.717563

**Authors:** Ping Liu, Lyuwei Wang, Shreya Basnet, Jianlin Cheng

## Abstract

Transcription factors (TFs) are central regulators of gene expression, and their selective recognition of genomic DNA underlies various biological processes. Experimental profiling of TF–DNA interactions using chromatin immunoprecipitation followed by sequencing (ChIP-seq) provides high-resolution maps of in vivo TF-DNA binding but remains costly, labor-intensive, and inherently low-throughput, limiting their scalability across different transcription factors, cell types, and regulatory conditions. Computational modeling therefore plays an essential role in inferring TF–DNA interactions at genome scale. However, most existing computational models rely solely on DNA sequence and chromatin features to predict TF–DNA binding, neglecting TF-specific protein information. This omission limits their ability to capture protein-dependent binding specificity. Here, we present TFBindFormer, a hybrid cross-attention transformer that explicitly integrates genomic DNA features with TF-specific representations derived from protein sequences and structures. By modeling protein-conditioned, position-specific TF–DNA interactions, TFBindFormer enables direct learning of molecular determinants underlying DNA recognition. Evaluated across hundreds of cell-type–specific TFs and hundreds of millions of genome-wide DNA bins, TFBindFormer consistently outperforms DNA-only baselines, achieving substantial gains in both area under precision-recall curve (AUPRC) and area under receiver operating characteristic curve (AUROC). Together, these results demonstrate that integrating TF and DNA features via cross-attention enables TFBindFormer to serve as an effective and scalable framework for large-scale TF–DNA binding prediction.

## Introduction

Transcription factors (TFs) are central regulators of gene expression, acting through the selective recognition of short, sequence-specific DNA elements—transcription factor binding sites (TFBSs)—to establish transcriptional programs that govern cellular identity, development, and function [1, 2]. Advances in high-throughput functional genomics assays, most notably chromatin immunoprecipitation followed by sequencing (ChIP-seq), have enabled genome-wide interrogation of TF–DNA interactions and substantially expanded our understanding of transcriptional regulation [3, 4]. However, despite these advances, experimentally derived binding maps remain sparse, highly condition-specific, and incomplete relative to the vast combinatorial space defined by TFs, cell types, developmental stages, and environmental contexts [5]. As a result, computational modeling has emerged as a critical and complementary strategy for systematically inferring TF–DNA binding landscapes at genomic scale.

Early computational models based on position weight matrices (PWMs) and motif scanning capture core DNA binding preferences, but are inherently limited in their ability to model contextual dependencies, cooperative binding, and higher-order regulatory grammar, owing to assumptions of positional independence and the lack of explicit modeling of interactions between binding motifs and surrounding sequence context [6–8]. These limitations motivated the development of more expressive computational frameworks capable of integrating broader sequence context and capturing complex regulatory interactions. The advent of deep learning, such as convolutional neural networks (CNNs), marked a major advance by enabling the direct learning of motif-like representations from raw genomic sequence, as demonstrated by DeepBind [9]. Subsequent large-scale studies further showed that architectural design choices—including network depth, pooling strategy, and receptive field size—play a critical role in determining predictive performance by governing how sequence context is encoded [10].

In parallel, multitask CNN frameworks such as DeepSEA extended this paradigm by jointly modeling diverse chromatin features, demonstrating that hierarchical representations and broader genomic context substantially improve TF-DNA binding prediction and enable single-nucleotide–resolution variant effect estimation [11]. Hybrid architectures combining CNNs with recurrent neural networks (RNNs), including DanQ and FactorNet, further enhanced predictive accuracy by capturing longer-range dependencies and incorporating additional regulatory signals, leading to improved cell-type–specific TF binding prediction [12, 13].

More recently, attention-based mechanisms have been integrated into transcription factor binding site prediction models to explicitly prioritize informative sequence regions, enabling more interpretable modeling of sequence determinants while preserving strong predictive performance [14]. A representative example is TBiNet, an attention-augmented CNN–BiLSTM architecture [15], which leverages attention to emphasize biologically relevant binding positions within DNA sequences. By selectively weighting core binding regions, TBiNet provides mechanistic insight into model predictions and has been shown to outperform earlier sequence-based approaches.

Despite these advances, the vast majority of existing TF-DNA binding predictors remain fundamentally DNA-centric, implicitly assuming that binding specificity is fully encoded in the genomic sequence alone. However, TF–DNA recognition is inherently a bidirectional process, shaped not only by DNA motifs and sequence context but also by TF protein sequence, structure, and biophysical properties [16, 17]. Recent work has begun to address this limitation by incorporating protein-level information into binding prediction. For example, TransBind integrates protein language model–derived embeddings with DNA features through the attention mechanism, enabling TF-specific modeling of genomic binding patterns and improving predictive performance[18]. Despite these advances, integrating TF protein features with genomic sequence context remains an area of ongoing research.

Recent advances in protein language modeling provide a compelling opportunity to incorporate protein-specific information into TF-DNA binding prediction, as they demonstrate that rich, transferable representations of protein sequences can be learned from large-scale datasets, enabling improved mechanistic insight into molecular recognition processes [19, 20]. At the same time, DNA sequence-based models have demonstrated that regulatory DNA context can be effectively encoded to support high-resolution TF-DNA binding prediction, as exemplified by attention-based architectures such as TBiNet. Together, these advances underscore the complementary representational strengths of modern protein and DNA encoders and provide a strong motivation for integrative models that explicitly couple transcription factor molecular features with genomic regulatory context.

Building on these advances, recent studies have begun to explore multimodal frameworks that jointly model transcription factor and DNA features, including cross-attention–based architectures that enable interaction between protein and genomic representations[21, 22]. These approaches further demonstrate the potential of explicitly modeling TF–DNA interactions to improve predictive performance and provide more interpretable insights into binding mechanisms. Cross-attention–based architectures, in particular, provide a principled framework for such integration by enabling fine-grained, position-aware interactions between protein and DNA representations while supporting structured, conditional information flow[23, 24].

Motivated by these considerations, we introduce TFBindFormer, a novel, bidirectional cross-attention transformer that integrates TF protein-specific embeddings with genomic DNA sequence features to model position-specific, protein-dependent DNA recognition. The model combines the modality-specific protein and DNA encoders with a hybrid cross-attention module that enables explicit residue-nucleotide interactions, producing TF-conditioned DNA representations for accurate TF–DNA binding prediction. By explicitly conditioning genomic predictions on TF features, TFBindFormer contributes to ongoing efforts in protein-aware modeling, providing an effective and scalable framework for modeling TF–DNA interactions across diverse TFs and regulatory contexts.

## Method

An overview of the architecture of TFBindFormer is shown in Figure 1. TFBindFormer is a hybrid bidirectional cross-attention model that integrates TF sequence and structure representations with genomic DNA sequence representations to predict TF–DNA binding across genome. The model integrates fixed, sequence- and structure-aware protein embeddings with DNA sequence features from a trainable DNA encoder and learns fine-grained residue-to-nucleotide interactions via a hybrid bidirectional cross-attention mechanism. The architecture comprises four major components: (i) a TF protein encoder block generating TF embeddings, (ii) a sequence-based DNA encoder block for genomic feature extraction, (iii) a hybrid cross-attention module that enables explicit residue–nucleotide interactions between protein and DNA representations, and (iv) a prediction head that aggregates the integrated features and maps them to a TF–DNA binding probability.

**Figure 1.**
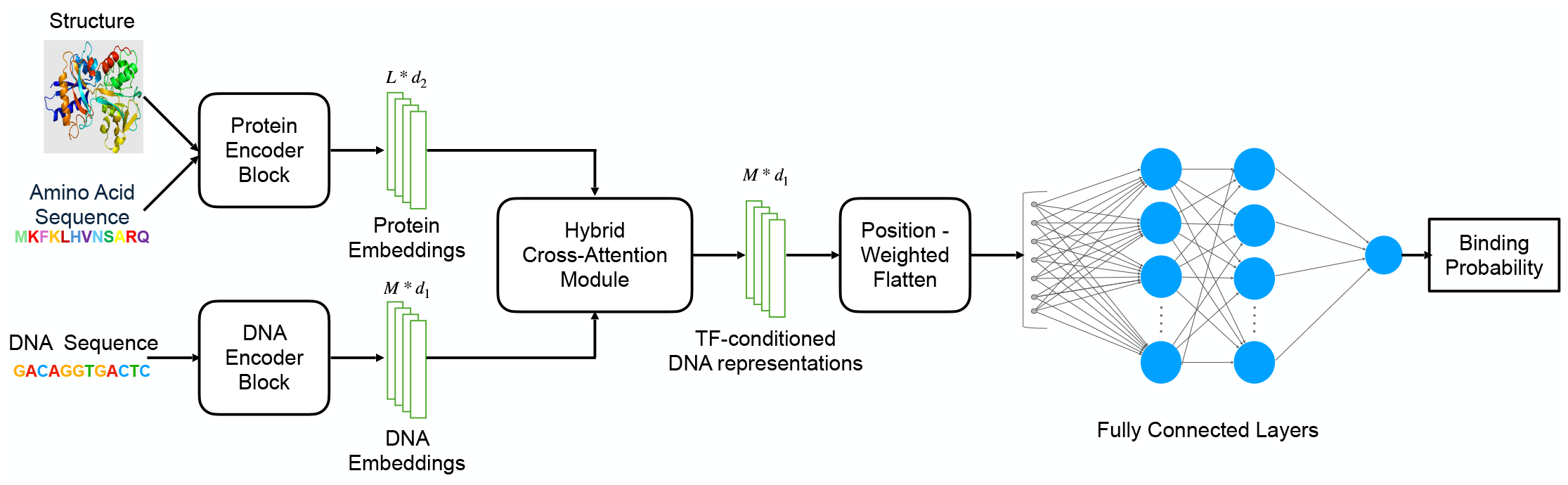
TFBindFormer architecture

### Protein Encoder Block

An overview of the protein encoder block is shown in Figure 2. The protein feature extraction module integrates primary sequence information with structure-derived geometric context to produce a unified residue-level representation. For each transcription factor (TF), its predicted tertiary structures obtained from the AlphaFold Protein Structure Database (AlphaFoldDB) [25, 26], was processed with Foldseek [27] to derive 3Di structural tokens. These tokens encode local protein backbone geometry and spatial relationships in a compact, sequence-like representation compatible with sequence-based modeling. The amino acid sequence together with its corresponding 3Di structural sequence is jointly provided as input to a pretrained protein language model - ProtST5 encoder [20], which produces contextualized residue-level embeddings from both sequence and structure modalities. The sequence- and structure-based protein embeddings are concatenated and linearly projected into a shared latent space of dimension *d*_2_, yielding a unified residue-level protein representation. All projection layers are trainable and shared across TFs.

**Figure 2.**
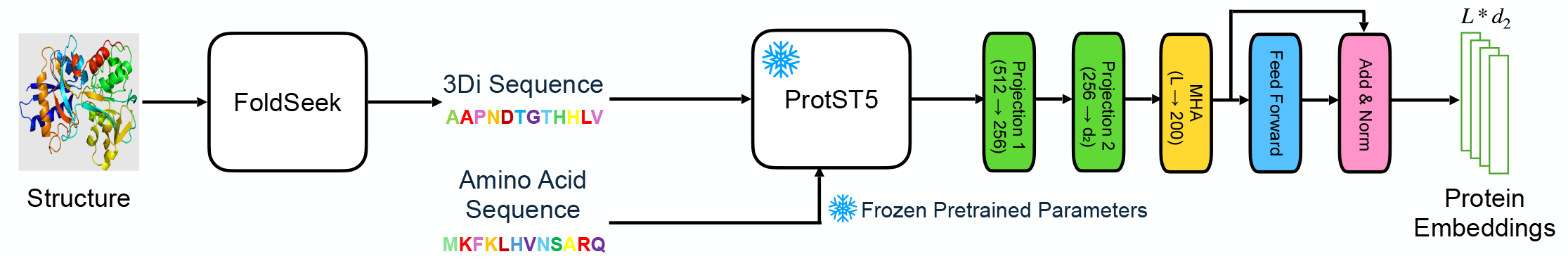
Protein encoder block

The resulting residue-level embedding matrix 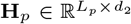 from the projection layers, where *L*_*p*_ denotes the length of the transcription factor protein sequence, is normalized to a fixed-length representation of *L* latent tokens using an attention-based reduction mechanism. Specifically, a set of *L* learned latent query vectors attends to the full residue sequence via multi-head attention (MHA), producing a compact global protein representation that aggregates information across the entire sequence.

The output of the MHA layer is further refined using a position-wise feed-forward network (FFN) followed by a residual connection and layer normalization:

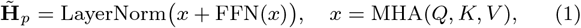

where FFN(·) consists of two linear transformations with a nonlinear activation in between. The resulting protein representation 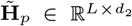 is used as an input to the hybrid cross-attention module.

### DNA Encoder Block

DNA features for each genomic bin are extracted using a TBiNet-inspired encoder [15], which combines convolutional motif detection, attention-based reweighting of informative sequence regions, and bidirectional recurrent modeling to capture both local and long-range regulatory dependencies. One-hot–encoded DNA inputs are first processed by convolutional layers with multiple receptive fields, followed by max pooling, to extract intermediate, motif-level sequence representations that capture recurring sequence patterns while reducing spatial resolution.

A lightweight attention mechanism is then applied to these representations to assign position-specific importance weights, enabling adaptive emphasis of discriminative motif instances and attenuating less informative background regions. The attention-weighted sequence features are subsequently processed by a bidirectional long short-term memory network (BiLSTM), which integrates contextual information across both upstream and downstream positions, yielding position-aware DNA embeddings that encode local motif features within their broader regulatory context.

To facilitate efficient and balanced interaction with the TF embeddings generated by the protein encoder in the downstream hybrid cross-attention module, DNA sequence representations are projected to a shared latent dimension *d*_1_ and subsequently resampled along the sequence axis to a fixed-length sequence of M tokens using nearest-neighbor interpolation. This resampling operation preserves relative positional ordering while providing a uniform input resolution independent of the original genomic bin length. The resulting M-token DNA embeddings serve as the DNA input to the hybrid cross-attention module, enabling explicit, position-specific integration with the TF representations.

### Hybrid Cross-Attention Module

An overview of the hybrid cross-attention module is shown in the upper part in Figure 3. The overall architecture of hybrid cross-attention module is composed of *n* repeated hybrid cross-attention blocks that explicitly model interactions between TF residues and DNA nucleotides, followed by a cross multi-head attention (Cross-MHA) layer, an addition and normalization layer, a feedfoward layer, and an addition and normalization layer to generate the final integrated TF-DNA binding features.

**Figure 3.**
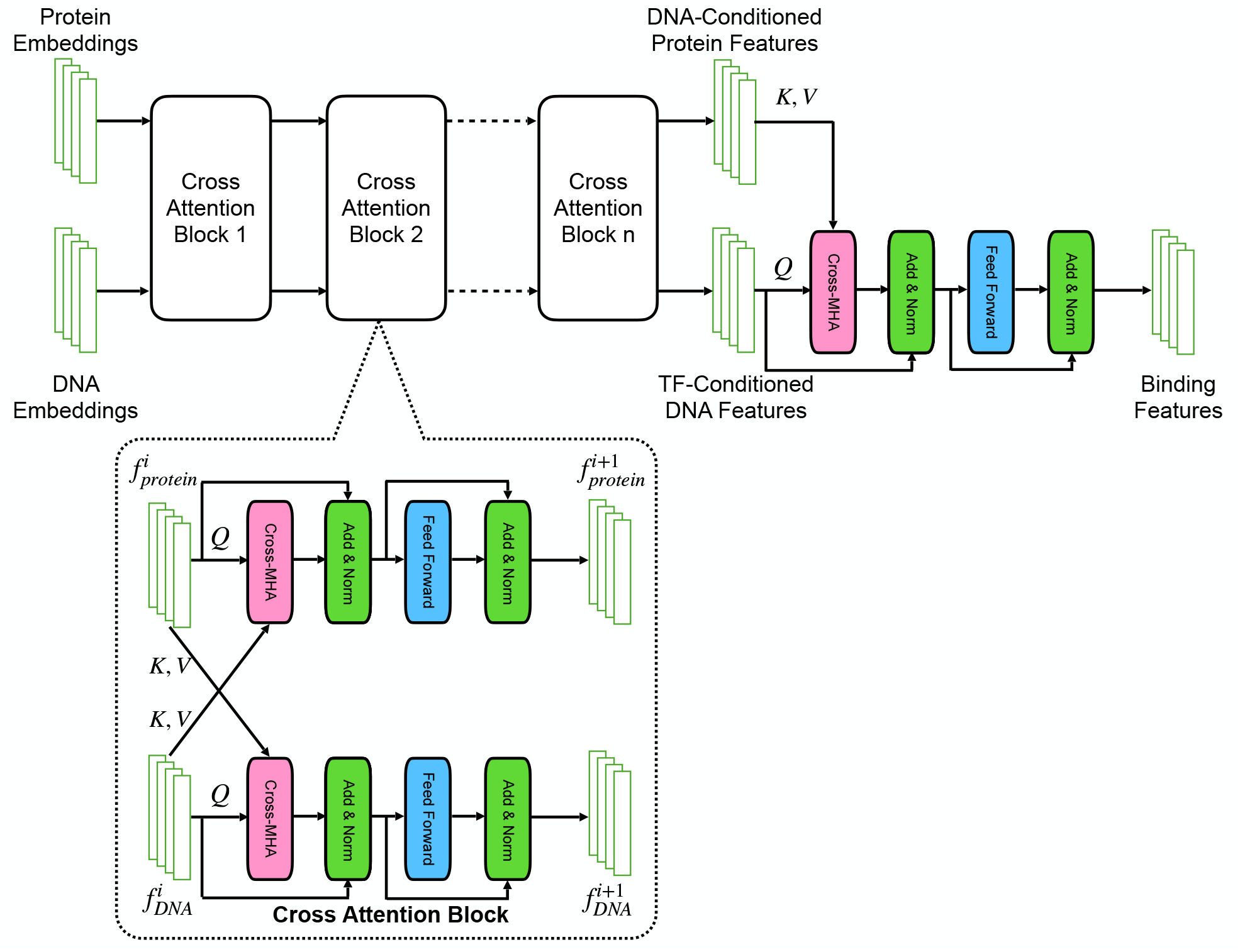
The hybrid cross-attention module.

Each hybrid cross-attention block operates on protein and DNA representations and performs reciprocal cross-attention, enabling direct information exchange between the two modalities. Through iterative stacking of these blocks, the model progressively refines both protein- and DNA-centric representations in a manner conditioned on their mutual compatibility.

Within a hybrid cross-attention block (the lower part in Figure 3), protein features attend to DNA features and DNA features simultaneously attend to protein features, allowing the model to jointly reason about sequence-specific DNA context and TF molecular properties. Protein-to-DNA cross-attention enables each protein residue embedding to selectively aggregate information from relevant DNA nucleotide positions, capturing potential contact patterns and preferences across the DNA sequence. Conversely, DNA-to-protein cross-attention allows each nucleotide position to integrate information from specific TF residues, facilitating residue-aware encoding of regulatory sequence features. This bidirectional formulation supports fine-grained, position-specific residue–nucleotide dependency modeling that cannot be captured by unidirectional or modality-isolated architectures.

Formally, protein features attend to DNA features according to:

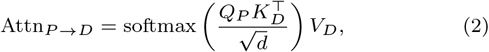

and, symmetrically, DNA features attend to protein features:

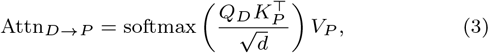

where *Q, K, V* denote the query, key, and value projections, and *d* is the attention head dimension. Each cross-attention operation is followed by residual connections, layer normalization, and position-wise feed-forward networks, ensuring stable optimization and enabling nonlinear transformation of the integrated representations.

The *n* cross-attention blocks employing fully bidirectional hybrid cross-attention to allow rich two-way interaction between protein and DNA modalities are followed by the final cross multi-head attention layer (Cross-MHA) adopting an asymmetric design in which only DNA positions attend to TF embeddings (Figure 3). This design choice reflects the predictive objective of TF–DNA binding and ensures that the final representation is explicitly DNA-centric yet conditioned on TF features. By suppressing a parallel protein output at the final stage, the encoder produces a single TF-conditioned DNA representation that directly encodes how each nucleotide position is modulated by TF features.

After the final cross-attention layer, TFBindFormer outputs TF-conditioned DNA representations of shape *M* × *d*_1_, where *M* denotes the fixed-length of the DNA token sequence. Each position in this representation captures contextualized TF–DNA interaction features that integrate local nucleotide information with TF sequence and structural context, serving as the input to the downstream prediction head for TF–DNA binding probability estimation.

### Prediction Head

The prediction head aggregates the TF-conditioned DNA representations produced by the hybrid cross-attention module and maps them to a TF-DNA binding probability (Figure 1). To obtain a sequence-level representation suitable for binding prediction, the resulting TF-conditioned DNA features of shape *M* × *d*_1_ are aggregated using a content-aware position-weighted pooling module, in which each nucleotide position is assigned a learned importance weight derived from its feature representation.

Specifically, given TF-conditioned DNA features:

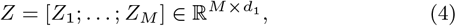

The flattened sequence representation is computed as follows:

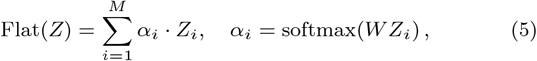

Where *W* denotes a learnable linear projection that produces scalar importance scores for each position. This content-aware flattening allows the model to adaptively emphasize biologically relevant regions, such as the central core binding region, while still incorporating contextual information from flanking sequences. The resulting flattened feature vector is then passed through a multilayer perceptron with normalization, nonlinear activation, and regularization to produce the final TF-DNA binding probability.

## Data and Experiment Setup

### Data Preparation

To construct a large-scale, uniformly processed dataset for TF–DNA binding prediction, we adopted the DeepSEA-style genome binning and labeling framework implemented in the build-deepsea-training-dataset pipeline[11]. Using the human reference genome (GRCh37/hg19), the genome was segmented into non-overlapping 200-bp bins, each of which was annotated for TF binding using high-quality ENCODE-derived ChIP-seq narrowPeak files of hundreds of TFs downloaded from the UCSC Genome Browser Database [28]. A bin was labeled positive for each TF if at least 50% of its length overlapped a ChIP-seq peak of the TF; otherwise, it was labeled as negative. This procedure produced a genome-wide binary label matrix spanning hundreds of cell-type–specific TFs and maintained compatibility with established DeepSEA-style TF binding benchmark.

For each labeled bin, we extracted a 1,000-bp genomic DNA sequence centered on the 200-bp target region, including ±400 bp of flanking context, to capture local regulatory context and potential higher-order motif interactions. Each sequence was encoded as a 1,000 × 4 one-hot matrix over the nucleotides A, C, G, and T.

Because our modeling objective focuses on learning sequence-specific protein–DNA interactions, we further refined the initial collection of 690 ChIP-seq experiments by restricting the dataset to TFs with intrinsic DNA-binding specificity. ChIP-seq experiments targeting histones, RNA polymerase, and other chromatin-associated proteins that associate with DNA indirectly were excluded. This biologically motivated filtering yielded a curated panel of 457 high-confidence cell-type–specific TF–DNA binding datasets. The resulting labeled data enables joint learning across diverse TF families and supports rigorous evaluation of models that integrate genomic DNA sequence with TF protein features.

To prevent information leakage and to assess unseen genomic loci, we employed chromosome-based data partitioning. The DNA bins of Chromosomes 4 and 7 were held out exclusively for validation, while chromosomes 8 and 9 served as an independent test set. All remaining chromosomes, excluding chromosome Y, were used for model training. Hyperparameters were tuned solely on the validation chromosomes and were never optimized using the test chromosomes.

### Dataset Statistics

The final dataset comprises the chromosome-partitioned training, validation, and test splits that differ substantially in size yet exhibit highly consistent TF-binding label distributions (Table 1). Each 200-bp genomic bin is annotated with 457 binary cell-type–specific TF labels, yielding a large-scale labeled dataset comprising approximately 2.38 billion TF–bin pairs (samples). Despite the severe class imbalance inherent to TF binding, all splits maintain a stable positive binding ratio of approximately 1% (train: 1.107%, validation: 0.980%, test: 1.055%), reflecting the sparse and localized nature of TF–DNA interactions across the genome. The large training split provides sufficient statistical coverage for scalable model learning, while chromosome-held-out validation and test splits enable unbiased hyperparameter tuning and rigorous assessment of unseen genomic regions.

**Table 1:**
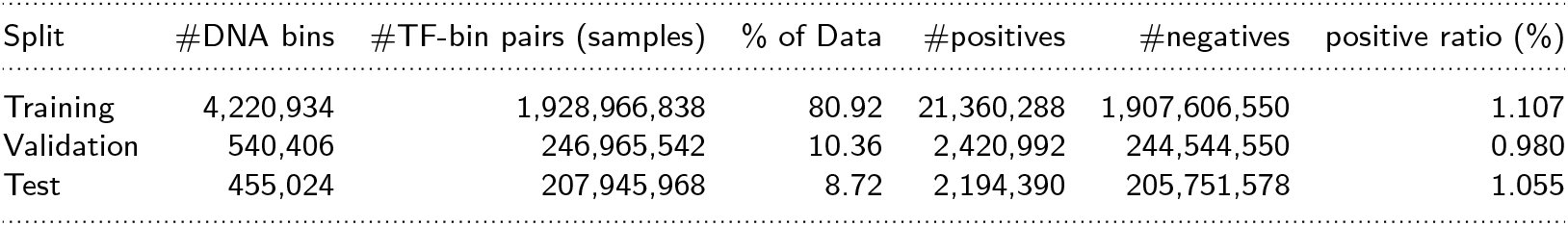
Statistics for Training, Validation and Test Data.

### Distribution of TF Binding Frequencies

To further assess the binding frequency of each TF in the large-scale labeled TF–DNA binding dataset, we examined the distribution of the number of positive genomic bins per TF across the training, validation, and test splits (Figure 4). Consistent with the label-level statistics reported in Table 1,violin plots with embedded boxplots reveal long upper tails and mean values substantially exceeding the median in all splits, indicating a strongly right-skewed, heavy-tailed distribution. This pattern reflects pronounced heterogeneity in TF binding activity, in which a small subset of TFs binds a disproportionately large number of genomic bins, whereas the majority exhibit much sparser binding across the human genome.

**Figure 4.**
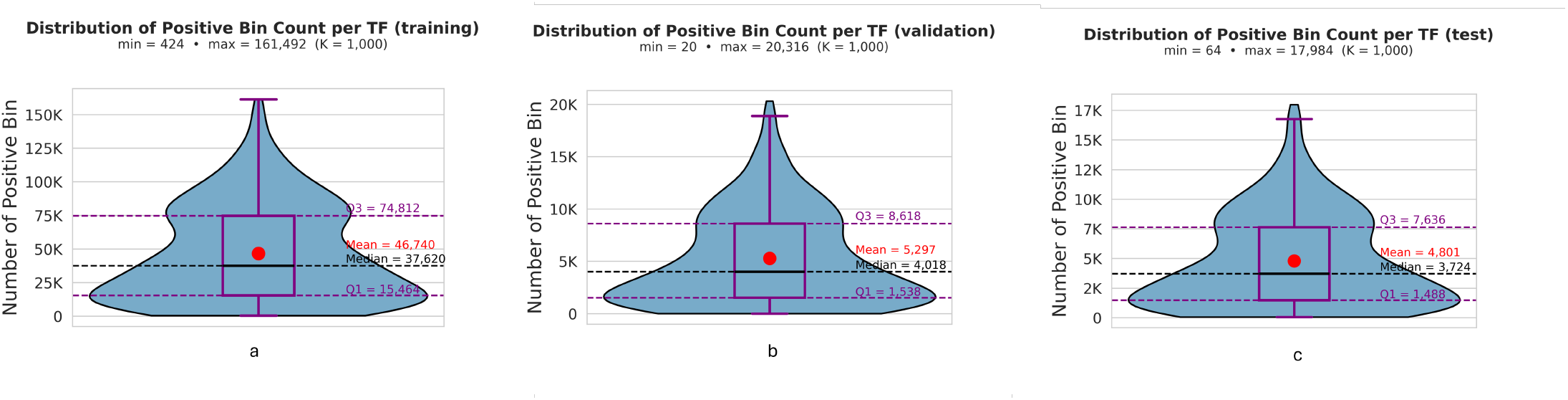
Distribution of the number of positive bins per TF in the training, validation, and test data splits. Violin plots with embedded boxplots show the distribution of positive genomic bins per cell-type-specific TF in the training (a), validation (b), and test (c) sets.

Across the training (Figure 4a), validation (Figure 4b), and test (Figure 4c) sets, per-TF positive bin counts span 424–161,492, 20–20,316, and 64–17,984, respectively. Despite these differences in absolute scale, all splits display a similar strongly right-skewed, heavy-tailed distribution, indicating that substantial heterogeneity in TF binding frequency is preserved across chromosome-partitioned splits.

### Experimental Setup

During the training experiments, several hyperparameters of TFBindFormer were set as follows: the numbers of DNA and protein tokens were set to *M* = 200 and *L* = 200, respectively, and the latent dimensions were set to *d*_1_ = *d*_2_ = 128.

The models were trained with a batch size of 1,024 for up to 20 epochs, corresponding to a total of 976,100 optimization steps. A linear warm-up phase of 1,000 steps was applied at the beginning of the training to stabilize optimization, and early stopping with patience of five epochs was used to mitigate overfitting. Data loading was parallelized using six worker processes.

Training was performed using the AdamW optimizer with a learning rate of 1×10^−4^ and a weight decay of 1×10^−5^. Following the warm-up phase, a cosine learning-rate decay scheduler was applied to improve convergence stability.

To address the severe class imbalance inherent in TF-DNA binding prediction, where the number of negative samples is substantially higher than the number of positive samples, different negative sampling fractions were used to balance the training and validation data. During training, the negative fraction was set to 0.15 to enrich positive samples. For validation, the negative fraction was increased to 0.5, while the test set retained all negative samples (negative fraction = 1.0) to reflect realistic genome-wide binding distributions.

## Results

### Overall predictive performance on held-out data

To comprehensively assess predictive performance on held-out data, we evaluated TFBindFormer using receiver operating characteristic (ROC) curve and precision–recall (PR) curve on the test set (Figure 5). The ROC curve exhibits a steep rise near the origin, yielding an area under the ROC curve (AUROC) of 0.956, which indicates excellent global discrimination between bound and unbound genomic binds across different decision thresholds on false positive rate.

**Figure 5.**
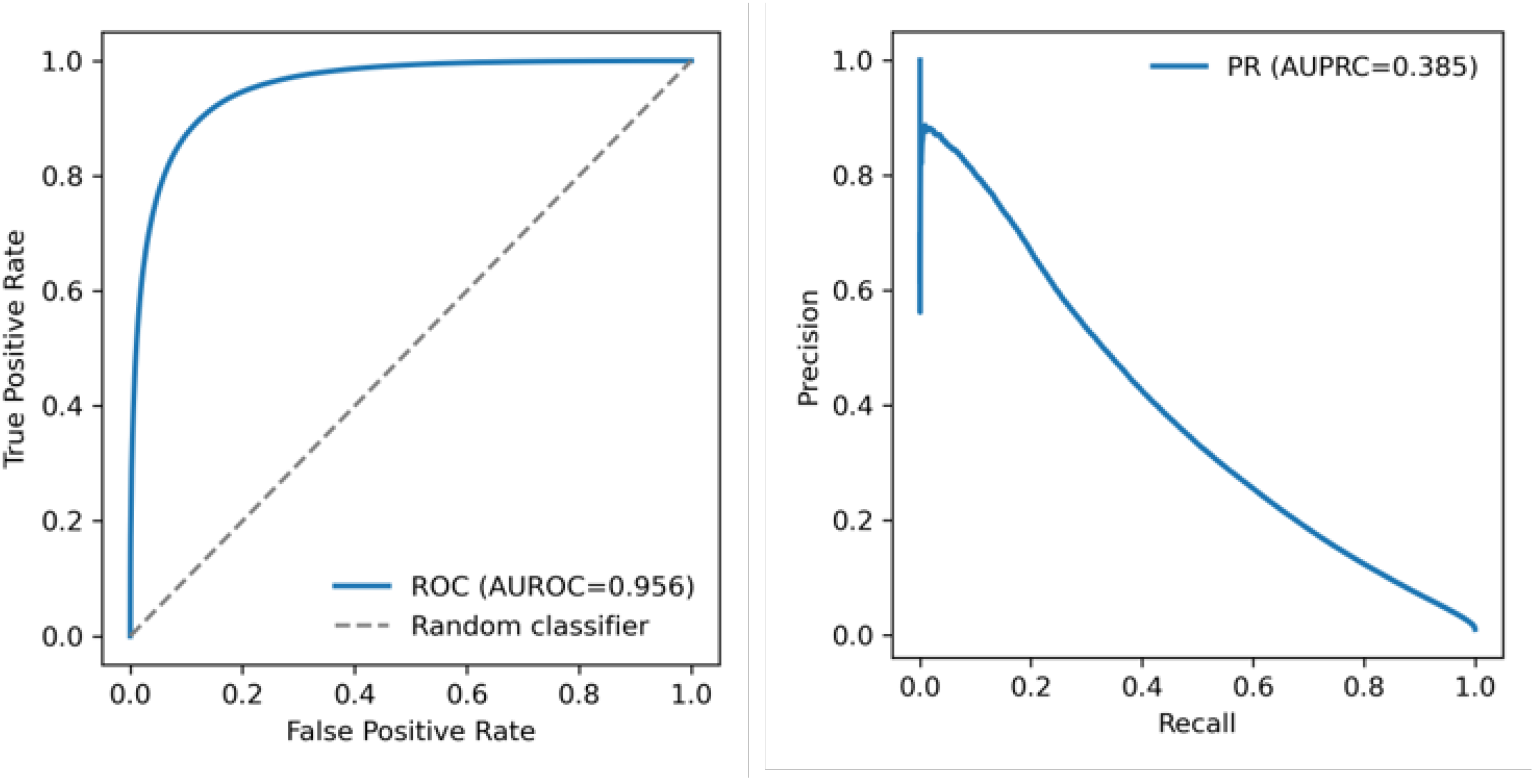
TF-DNA binding prediction performance of TFBindFormer on the test data. Left: the ROC curve of TFBindFormer; Right: PRC curve.

Due to the highly imbalanced nature of TF-DNA binding data, i.e., only about 1% of bins being positive, precision-recall analysis provides a more stringent evaluation of positive prediction quality. TFBindFormer achieved an area under PR curve (AUPRC) of 0.385, corresponding to a 36-fold improvement over the random baseline, defined by the positive binding site prevalence in the test set (1.055%; Table 1), and demonstrating substantial enrichment of true binding events among high-scoring predictions. Precision is highest at low recall and decreases smoothly as recall increases, indicating that the model prioritizes a subset of high-confidence binding sites while maintaining an informative ranking across decision thresholds.

### Per-transcription factor performance analysis

While aggregate evaluation demonstrates strong overall predictive performance, we further conducted a fine-grained analysis at the level of individual transcription factors to assess model robustness across diverse regulatory contexts. The benchmark includes 108 transcription factors (TFs), each was evaluated independently using aggregated predictions across all available cell types.

As shown in (Figure 6a), AUROC values remain consistently high across TFs, indicating that TFBindFormer achieves stable global discrimination between bound and unbound genomic regions for a wide range of transcription factors. In contrast, AUPRC values show greater variability across TFs (Figure 6b) reflecting differences in task difficulty and class imbalance. For example, TFs such as CTCF, Znf143, MafF, MafK, and ELK1 achieve notably higher performance (Table 2), consistent with their well-characterized DNA-binding motifs.

**Table 2:**
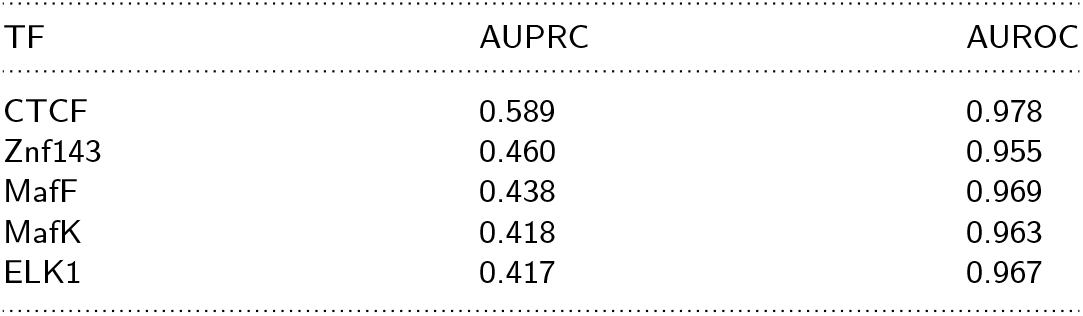
Top five transcription factors ranked by AUPRC on the held-out test set.

| TF | AUPRC | AUROC |
| --- | --- | --- |
| CTCF | 0.589 | 0.978 |
| Znf143 | 0.460 | 0.955 |
| MafF | 0.438 | 0.969 |
| MafK | 0.418 | 0.963 |
| ELK1 | 0.417 | 0.967 |

**Figure 6.**
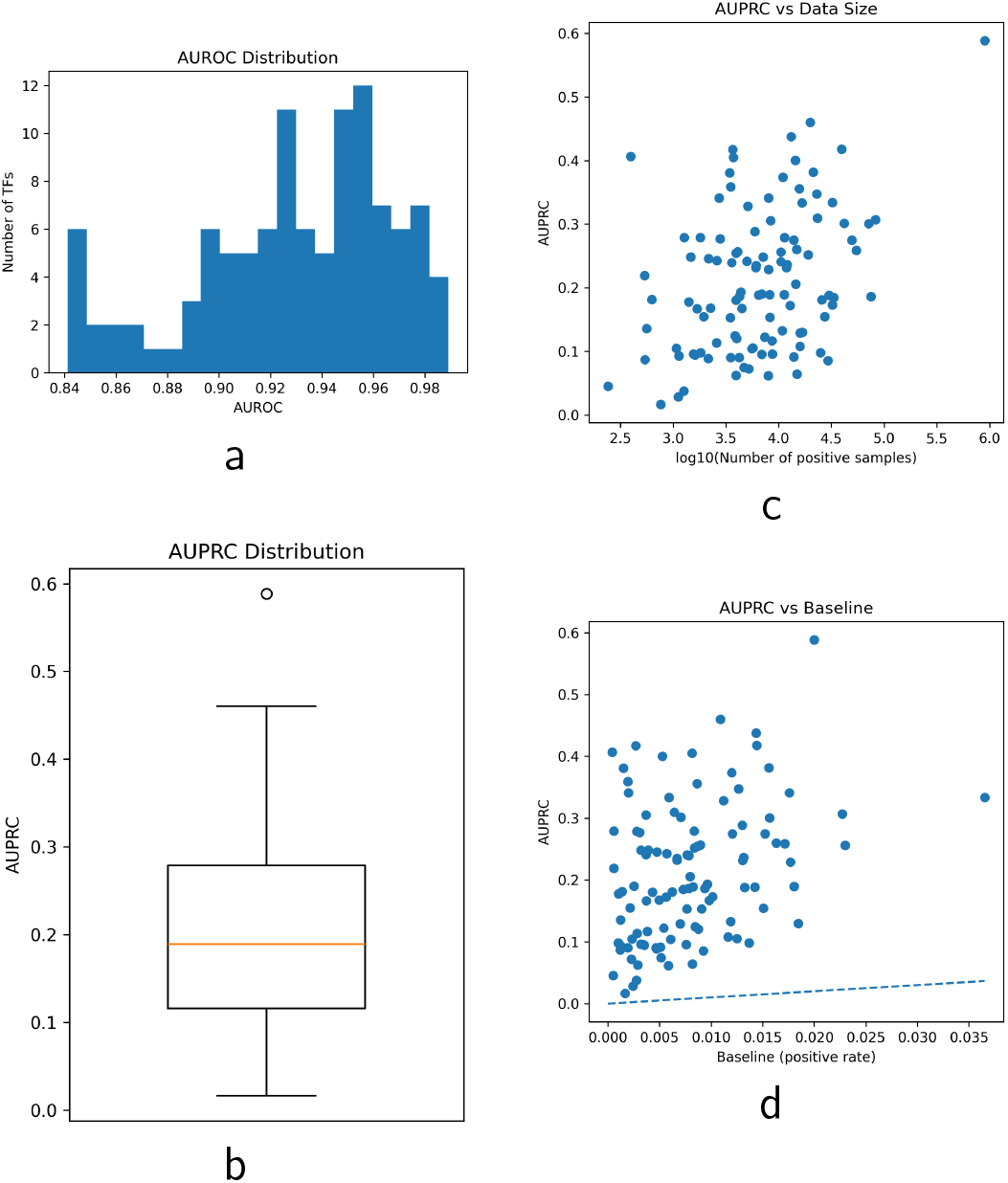
Per-transcription factor performance analysis. (a) Distribution of AUROC across TFs, showing consistently high discrimination. (b) Distribution of AUPRC, indicating variability in prediction performance across TFs. (c) Relationship between AUPRC and the number of positive samples per TF (log scale), demonstrating dependence on data availability. (d) AUPRC versus baseline positive rate, showing consistent improvement over random expectation across TFs.

We further examined the relationship between predictive performance and data availability of positive examples. As shown in (Figure 6c), AUPRC increases with the number of positive samples per TF, suggesting that positive example abundance contributes to improved performance. However, considerable variability remains among TFs with similar sample sizes, indicating that intrinsic biological properties, such as motif strength and context-dependent binding, also play an important role in determining prediction difficulty.

To account for class imbalance, we compared AUPRC against the baseline positive rate for each TF. TFBindFormer consistently achieves performance well above random expectation across TFs (Figure 6d), demonstrating that the model captures meaningful regulatory signals beyond simple data-driven effects.

Notably, TFs with well-defined sequence motifs tend to exhibit higher predictive performance, whereas TFs with more context-dependent or cooperative binding patterns remain comparatively challenging. These results indicate that TFBindFormer effectively models sequence-driven binding specificity while also highlighting the inherent complexity of transcriptional regulation.

### Comparison with Existing Deep Learning Methods

We compared overall predictive performance of TFBindFormer against four existing representative deep learning TF-DNA binding prediction models, including DeepSEA [11], DanQ [12], TBiNet [15], and EPBDXDNABERT-2 [29] on the held-out test set (Table 3). For a fair comparison, all models except EPBDXDNABERT-2 were trained using the same training and validation data splits and blindly evaluated on identical held-out test data. For EPBDXDNABERT-2, as certain components of the model implementation are not publicly available, we were unable to retrain the model under our experimental setting. Therefore, we report the performance metrics as provided in the original publication. The PR and ROC curves for all other models are shown in Figure 7.

**Table 3:** Performance of TFBindFormer and other deep learning methods on the held-out test set.

| Model | AUPRC | AUROC |
| --- | --- | --- |
| DeepSEA | 0.272 | 0.922 |
| DanQ | 0.290 | 0.928 |
| TBiNet | 0.310 | 0.938 |
| EPBDXDNABERT-2 | 0.326 | 0.949 |
| TFBindFormer | **0.385** | **0.956** |

**Figure 7.**
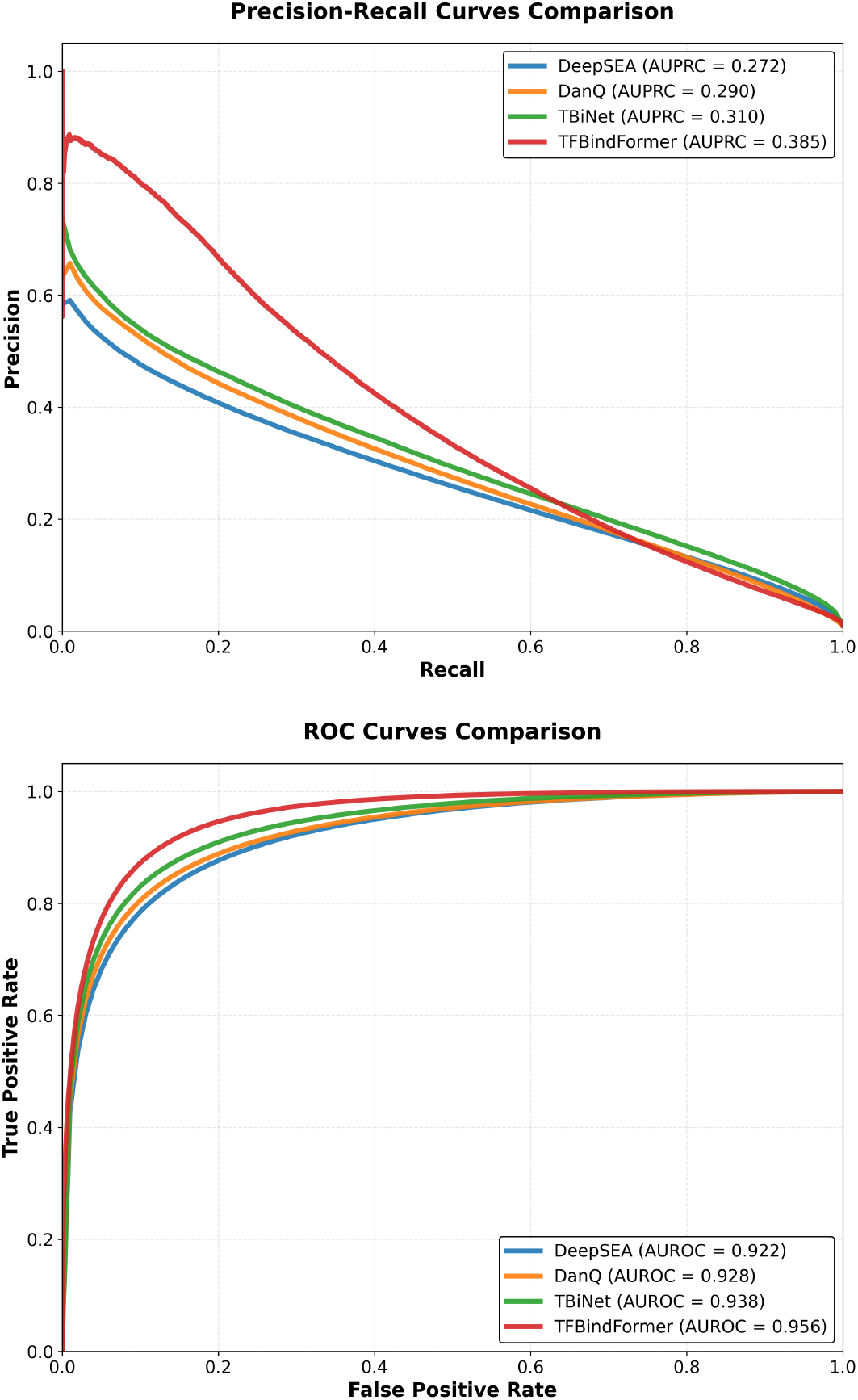
The Precision-Recall (PR) curves (top) and ROC curves (bottom) of all methods on the test data. TFBindFormer performed best according to both metrics, with particularly substantial improvements in terms of the PR curve.

TFBindFormer achieved the strongest overall performance in terms of both AUPRC and AUROC. In particular, it attained an AUPRC of 0.385, corresponding to relative improvements of 41.5%, 32.8%, 24.2%, and 18.1% over DeepSEA, DanQ, TBiNet, and EPBDXDNABERT-2, respectively. These gains indicate substantially improved enrichment of true TF binding events under the severe class imbalance condition and reflect the model’s enhanced ability to prioritize biologically relevant binding sites among high-confidence predictions.

TFBindFormer also achieved the highest AUROC of 0.956, exceeding DeepSEA (0.922), DanQ (0.928), TBiNet (0.938), and EPBDXDNABERT-2(0.949), demonstrating consistent gains in global discrimination between bound and unbound genomic bins.

Notably, while TFBindFormer adopts a TBiNet-style DNA encoder block, it significantly improves prediction performance by incorporating TF-specific protein representations and integrating TF and DNA features via explicit TF-DNA cross-attention. The consistent performance improvements over TBiNet therefore highlight the added value of explicitly modeling TF-DNA interactions beyond DNA sequence information alone.

### Ablation Study

To quantify the contribution of protein sequence and structural information to TF-DNA binding prediction, we performed an ablation analysis in which amino acid sequence embeddings or 3Di structural representations were selectively removed from TFBindFormer (Figure 8). Across both evaluation metrics (AUROC and AUPRC), the full model consistently achieved the strongest performance. AUROC was slightly reduced under both ablations, whereas the removal of amino-acid sequence information resulted in a pronounced degradation in AUPRC (−0.013), indicating a substantial loss in precision-recall enrichment under the severe class imbalance. In contrast, removing 3Di structural information led to a smaller but consistent reduction in AUPRC (−0.005), suggesting that structural cues provide complementary, though secondary, contributions to binding discrimination.

**Figure 8.**
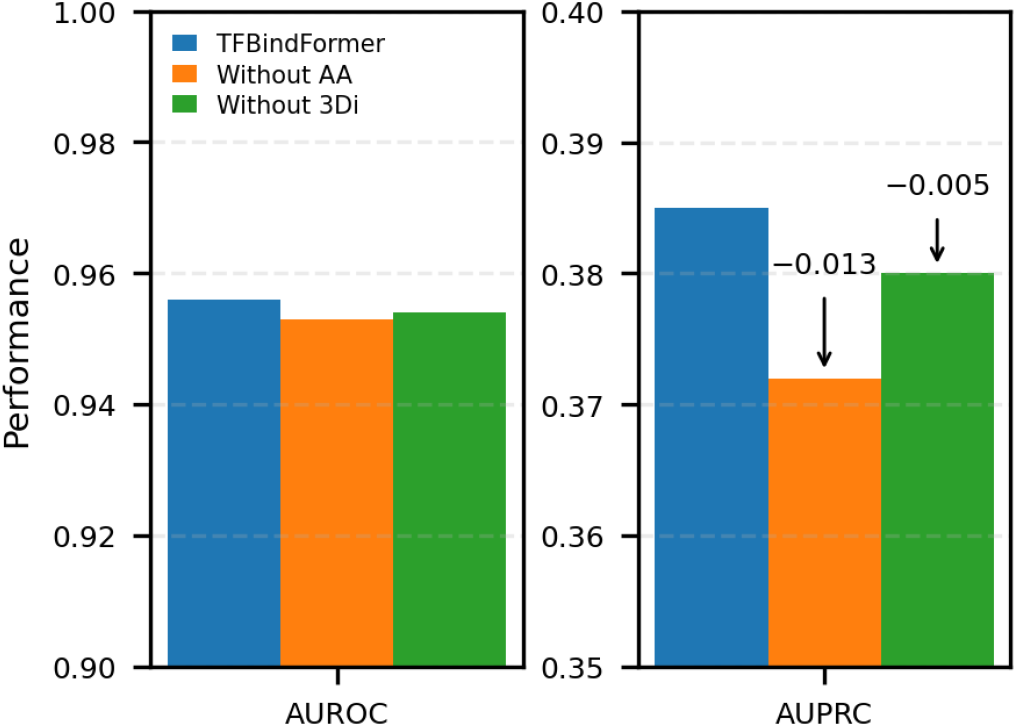
Ablation analysis of TFBindFormer. Performance comparison between the full model and variants with amino acid sequence (AA) or 3Di structural representations removed. AUROC is shown in the left panel, while AUPRC is displayed in the right panel. Removal of amino acid (AA) information leads to the largest reduction in AUPRC, whereas removing 3Di structural features also causes some performance loss.

Together, these results demonstrate that protein sequence constitutes the dominant signal driving TF-conditioned DNA recognition, while structure-derived information further refines prediction accuracy.

### Interpretation of TF–DNA Recognition via Attention Scores

To assess how TFBindFormer encodes TF-DNA binding specificity, we examined final-layer TF→DNA cross-attention patterns for a representative TF (CTCF) in the GM12878 cell line. Two representative genomic DNA bins were selected, one experimentally annotated as bound (positive) and another unbound (negative), for which the model produced highly confident but opposing predictions, with binding probabilities of 0.941536 and 0.000075 for the two bins, respectively.

From the final hybrid cross-attention layer, we extracted the CTCF→DNA attention matrix **A**^(*L*)^ ∈ R^200×200^, where each entry denotes the attention score from a CTCF token to a DNA token, averaged across attention heads. Averaging attention across CTCF tokens for each DNA token yields a DNA-level attention profile that explain how the prediction for the two bins were made. For the bound bin, the mean attention displays a sharp peak within the central region (DNA tokens 80–120) that contains the TF binding motif and rapidly decays outside it, whereas the unbound bin that does not contain a TF binding motif exhibits a flat, low-amplitude profile, with attention values remaining below 0.02 across all positions (Figure 9).

**Figure 9.**
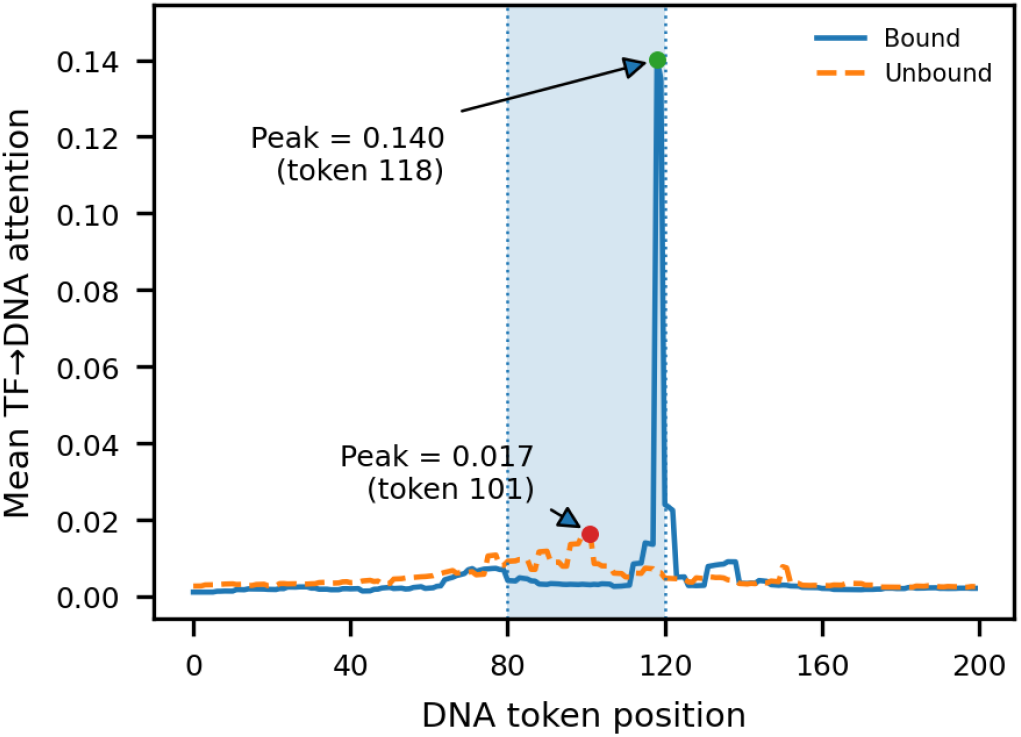
TF→DNA attention profiles for CTCF-bound and unbound genomic bins. TF→DNA cross-attention weights from the final layer were averaged across TF tokens to obtain DNA-centric attention profiles across DNA tokens. The blue curve (Bound) corresponds to a representative CTCF-bound bin, while the dashed orange curve (Unbound) represents a CTCF-unbound bin. The shaded region marks the central region (DNA tokens 80–120, corresponding to the central 200-nucleotide region) that contains the CTCF binding site in the bound bin.

Together, these results demonstrate that high-confidence CTCF binding predictions are associated with strong final-layer attention scores concentrated in the motif-enriched core region, whereas non-binding bins lack both elevated attention magnitude and spatial localization. This establishes a quantitative and interpretable link between the model’s learned attention structure and its discrimination of bound versus unbound DNA bins.

## Conclusion

We present TFBindFormer, a hybrid cross-attention framework that integrates transcription factor (TF) sequence and structure representations with genomic DNA sequence context for TF–DNA binding prediction. By combining protein embeddings with compressed DNA representations via cross-attention, the model enables explicit, position-specific modeling of protein–DNA interactions beyond DNA sequence-only approaches. Across genome-wide benchmarks, TFBindFormer consistently outperformed existing deep learning models, including DeepSEA, DanQ, TBiNet and EPBDXDNABERT-2, achieving substantial improvements in both AUROC and AUPRC under severe class imbalance. Notably, gains in AUPRC indicate improved enrichment of true positive binding sites among high-confidence predictions, a critical requirement for large-scale regulatory discovery.

Moreover, the ablation analyses show that these gains may be driven primarily by protein amino-acid sequence information and secondarily by protein structure information. Consistent with these findings, the cross-attention visualizations of some examples reveal biologically meaningful patterns, with attention concentrated over motif-enriched central binding regions in bound DNA sequences and diffuse, attenuated interactions in unbound regions. Together, these results demonstrate that protein-aware hybrid cross-attention modeling improves accuracy, robustness, and interpretability for TF–DNA binding prediction, providing an effective and scalable framework for integrating protein information into regulatory genomics.

## Abbreviations

TF: Transcription Factor
AUROC: area under receiver operating characteristic curve
AUPRC: area under precision-recall curve
MHA: multi-head attention

## Data availability

The data underlying this article are available in https://doi.org/10.5281/zenodo.18362288. The source code of this study are available at https://github.com/BioinfoMachineLearning/TFBindFormer

## Author contributions statement

J.C. conceived the experiment; P.L., L.W., S.B. conducted the experiment and collected data; P.L., L.W., S.B., and J.C. analysed the results; P.L., L.W., S.B., and J.C. wrote and reviewed the manuscript.

